# Global siRNA Screen Reveals Critical Human Host Factors of SARS-CoV-2 Multicycle Replication

**DOI:** 10.1101/2024.07.10.602835

**Authors:** Xin Yin, Yuan Pu, Shuofeng Yuan, Lars Pache, Christopher Churas, Stuart Weston, Laura Riva, Lacy M. Simons, William J. Cisneros, Thomas Clausen, Paul D. De Jesus, Ha Na Kim, Daniel Fuentes, John Whitelock, Jeffrey Esko, Megan Lord, Ignacio Mena, Adolfo García-Sastre, Judd F. Hultquist, Matthew B. Frieman, Trey Ideker, Dexter Pratt, Laura Martin-Sancho, Sumit K Chanda

## Abstract

Defining the subset of cellular factors governing SARS-CoV-2 replication can provide critical insights into viral pathogenesis and identify targets for host-directed antiviral therapies. While a number of genetic screens have previously reported SARS-CoV-2 host dependency factors, these approaches relied on utilizing pooled genome-scale CRISPR libraries, which are biased towards the discovery of host proteins impacting early stages of viral replication. To identify host factors involved throughout the SARS-CoV-2 infectious cycle, we conducted an arrayed genome-scale siRNA screen. Resulting data were integrated with published datasets to reveal pathways supported by orthogonal datasets, including transcriptional regulation, epigenetic modifications, and MAPK signalling. The identified proviral host factors were mapped into the SARS-CoV-2 infectious cycle, including 27 proteins that were determined to impact assembly and release. Additionally, a subset of proteins were tested across other coronaviruses revealing 17 potential pan-coronavirus targets. Further studies illuminated a role for the heparan sulfate proteoglycan perlecan in SARS-CoV-2 viral entry, and found that inhibition of the non-canonical NF-kB pathway through targeting of BIRC2 restricts SARS-CoV-2 replication both *in vitro* and *in vivo*. These studies provide critical insight into the landscape of virus-host interactions driving SARS-CoV-2 replication as well as valuable targets for host-directed antivirals.

## INTRODUCTION

As of May 2024, severe acute respiratory syndrome coronavirus 2 (SARS-CoV-2), the causative agent of COVID-19, has infected more than 775 million people worldwide and led to over 7 million deaths according to the World Health Organization (WHO). In the last 21 years, other coronaviruses have caused zoonotic outbreaks of severe viral respiratory illness in the human population. These include SARS-CoV-1, which was first reported in 2003 and has caused over 8,000 infections with a mortality rate of 9.5%^1^, and MERS, which was initially reported in 2012 and responsible for over 2,500 infections with a 34.4% fatality rate^2^. Four years after the SARS-CoV-2 pandemic was declared and despite available therapeutics and vaccines, the virus still remains a global health threat due to vaccine hesitancy, limited rollout of vaccines in certain demographic areas, and the surge of variants with increased immune evasion, replicative fitness, and transmission^3,4^. Elucidating host-pathogen interactions that are critical for SARS-CoV-2 replication can facilitate the understanding of SARS-CoV-2 biology and the development of host-directed antivirals that could benefit from broad-spectrum activities and reduced viral resistance^5,6^.

SARS-CoV-2 belongs to the family of enveloped viruses known as *Coronaviridae*^7^, which are enveloped, positive strand RNA viruses^8^. Virions are spherical and decorated with Spike (S) glycoproteins, which mediate receptor binding to facilitate viral entry^9^. Upon internalization, the viral RNA is released into the cytoplasm and transcribed into viral proteins^10^. These include structural proteins S, Envelope (E), Nucleocapsid (N), and Membrane (M) proteins, as well as 16 non-structural and 9 accessory proteins that are important for viral replication, innate immune evasion, and pathogenesis^11,12^. Coronaviruses induce the formation of double-membrane vesicles to promote the replication and transcription of their genomes^13^. Newly synthesized genomic RNAs are incorporated into virions and, following budding, infectious viruses are released from the host cell. Throughout their entire replication cycle, coronaviruses co-opt host factors that provide essential activities, including the cellular receptor ACE2 that is required for viral entry^14^. Previous CRISPR functional genetic screens have illuminated host factors and cellular pathways that are required for replication of SARS-CoV-2 and other coronaviruses^15–25^. However, these CRISPR screens were conducted in a pooled format, biasing them to the identification of host factors affecting initial stages of viral replication. Therefore, the host factor requirements for SARS-CoV-2 egress and budding remain poorly characterized.

Here, we report findings of an arrayed genome-wide siRNA screen to identify host factors involved throughout the entire SARS-CoV-2 infectious cycle. These factors were subsequently validated using targeted CRISPR-Cas9 technologies and integrated with previously reported OMICs, including functional genetics and proteomics, to reveal transcriptional control, epigenetic regulation and MAPK signalling as pathways implicated in SARS-CoV-2 replication with support from multiple studies. Proviral host factors were then mapped for their ability to support distinct stages of the SARS-CoV-2 infectious cycle, e.g., entry, viral RNA replication/translation, or egress, and we found that the majority of host factors impact replication or egress. In addition, we identified 17 potential pan-coronavirus host factors, including perlecan, which was found to facilitate viral entry and was determined as a direct interactor of SARS-CoV-2 S protein. Small molecules targeting the proviral factor Baculoviral IAP Repeat Containing 2 (BIRC2) were found to inhibit SARS-CoV-2 infection in a dose-dependent manner. The proviral effects of BIRC2 on SARS-CoV-2 growth were further confirmed *in vivo* by treating infected mice with a BIRC2 inhibitor. Overall, this study provides new insights into host factors required for the entire SARS-CoV-2 replication cycle, including late stages, and identifies host-targeting inhibitors that can serve as the basis for new anti-SARS-CoV-2 therapies.

## RESULTS

### Genome-wide screen identifies host factors involved in SARS-CoV-2 replication

The systematic identification of cellular factors that either support or restrict viral replication can provide valuable insights into SARS-CoV-2 biology, pathogenesis, and identify new antiviral targets. To uncover host factors involved in SARS-CoV-2 replication, we conducted a genome-wide siRNA screen in human Caco-2 cells challenged with USA-WA1/2020, the first SARS-CoV-2 US isolate (**Figure 1A**). This colorectal adenocarcinoma cell line was selected for the screen because the intestinal epithelium is a target for SARS-CoV-2^26,27^ and these cells endogenously express ACE2 and TMPRSS2, rendering them permissive to SARS-CoV-2 infection^14^. Furthermore, the siRNA knockdown efficiency is higher in Caco-2 cells compared to other SARS-CoV-2 permissive cell types such as Calu3. Cells were transfected with individually arrayed siRNAs, infected with SARS-CoV-2 for 48 h, immunostained for SARS-CoV-2 N protein, stained with DAPI, and then subjected to high content microscopy (**Figure 1A**). The impact of each individual gene knockdown on viral replication (% infected cells) was quantified based on DAPI^+^ events (number of cells) and SARS-CoV-2 N^+^ events (number of infected cells), and then normalized to the median % infection of each plate. Non-targeting, scramble siRNAs were included on each plate as negative controls, and siRNAs targeting SARS-CoV-2 entry factors ACE2 and TMPRSS2 were included as positive controls (**Figure S1A**). Screens were conducted in duplicate and showed good reproducibility with a Pearson correlation coefficient (r) = 0.66 (**Figure S1B**). Primary screening data were subjected to an analysis pipeline to identify siRNAs that affect viral replication (ranked based on Z-score) without impacting cell viability (cell count at least 70% of scramble control). Using these criteria, we identified 253 proviral host factors (including 222 with Z-scores < −2 in both replicates, and 31 with Z-score < −2 in replicate 1 and < −1.5 in replicate 2) (**Figure 1B**, green). Additionally, we identified 81 factors that restricted viral replication (Z-score > 1.5 in both replicates), including CCND3, which we previously identified as a restriction factor for SARS-CoV-2^28^ (**Figure 1B**, red). Findings are summarized in **Table S1**. Reactome and gene ontology (GO) analyses of proviral factors revealed enrichment in intracellular protein transport (LogP=-3.5398), proteosome-mediated ubiquitin process (LogP=-3.1010), and cell junction organization (LogP=-2.7385), among the top 10 enriched terms (**Figure 1C**, left). Antiviral factors were enriched in protein phosphorylation (LogP=-8.1590), JAK-STAT signalling (LogP=-4.0693), and demethylation (LogP=-3.7072), amongst others (**Figure 1C**, right). Gene membership to these terms is included in **Table S1**. Host factors identified in the primary screen were subjected to a subsequent round of siRNA validation using four individually arrayed siRNAs per gene to minimize off-target effects. Here, 125 cellular factors were confirmed to affect the replication of SARS-CoV-2 with 2 or more siRNAs (**Figure 1D**) and their expression was verified across different relevant cell types^29^, including primary mucocilliated epithelial cells, which are a known target of SARS-CoV-2 (**Figure S2A**). We also further validated a subset of 12 factors using CRISPR-Cas9 knockout in the human lung cell line Calu-3 (**Figure 1E**). Combined, these data provide a list of validated host factors across different cell types that are involved in SARS-CoV-2 replication.

**Figure 1.**
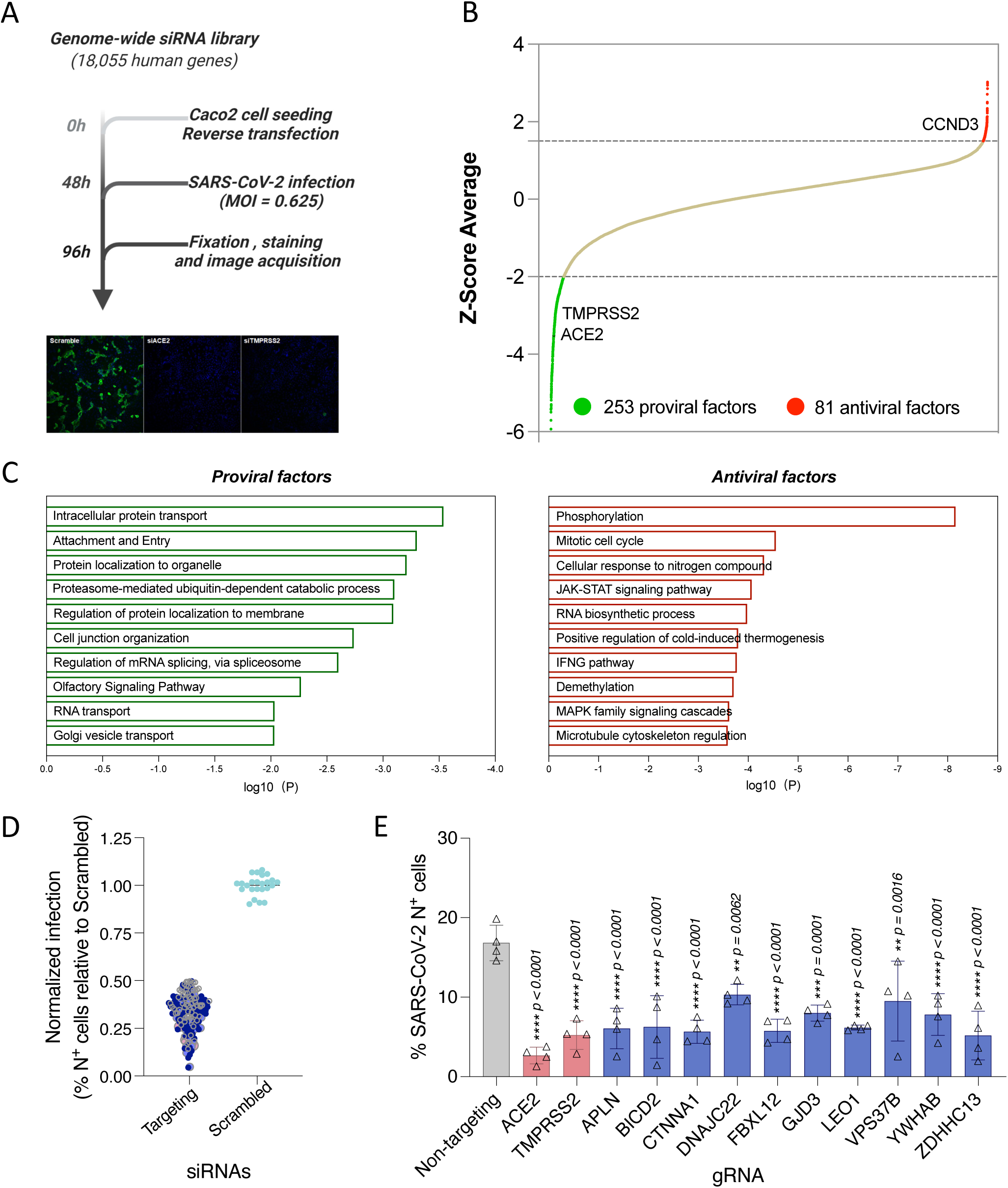
Genome-wide siRNA screen identifies host factors involved in SARS-CoV-2 replication. (A) Schematic representation of the genome-wide screen to identify human host factors that affect SARS-CoV-2 replication. (B) Ranked SARS-CoV-2 infectivity Z-scores from the genome-wide siRNA screen. Dashed lines illustrate cut-offs for hit calling strategy: Z-score ≤ −2 indicates proviral factors (green), Z-score ≥ 1.5 indicates antiviral factors (red). Controls are shown (e.g., siACE2, positive). (C) Functional enrichment analysis of identified proviral (*left-green*) and antiviral (*right-red*) host factors. (D) Deconvolution plot showing proviral host factors validated with one siRNA (grey), two siRNAs (dark blue), three siRNAs (light blue) and four siRNAs (pink). (E) Calu-3 cells treated with indicated gRNAs were infected with SARS-CoV-2 (MOI = 0.75) for 48 h prior to immunostaining for viral N protein. Shown is quantification of the normalized infection (% of SARS-CoV-2 N^+^ cells) relative to parental cells. Data show mean ± SD from one representative experiment in quadruplicate (n=4) of two independent experiments. Significance was calculated using one-way ANOVA with Dunnett’s post-hoc test.

### Network integration reveals transcriptional control, epigenetic modifications, and MAPK signalling as relevant networks implicated in SARS-CoV-2 replication

SARS-CoV-2 relies on a number of cellular proteins to complete its replication cycle, from surface receptors for viral entry to vesicle transport and sorting proteins for viral trafficking and release^30^. Conversely, in response to infection, the cell activates an antiviral program to clear infection^28^. A network integration model was generated to identify the interactomes and networks that the SARS-CoV-2 proviral and antiviral factors identified in our primary screen belong to and thereby gain a better understanding of their role in viral replication. First, we conducted a supervised network propagation by creating a grid that included the siRNA screening hits and their high confidence interactors as determined by the STRING database (*see Methods*). To put the host factors that we identified in context of previously identified SARS-CoV-2 host factors and highlight more confidence networks and host factors, we leveraged the first two reported SARS-CoV-2 functional genetic screens^15,16^, as well as the first two reported SARS-CoV-2 interactome and a phosphoproteomics datasets^31–33^. These datasets were integrated with the genetic screen data generated in this study and community detection algorithms were applied to identify densely interconnected clusters of factors that show significant membership in biological processes (**Figure 2**; *see Methods*). The resulting hierarchical ontology network revealed enrichment in metabolic pathways (p value = 2.83E-23) (**Figure S2B**), which were previously reported to affect viral replication by controlling cellular energy levels^34^, as well as enrichment in vesicle transport (p value = 7.62E-9). The vesicle transport cluster included factors such as Clathrin heavy chain 1 (*CLTC*), important for entry of several RNA viruses^35^, and the vacuolar protein sorting associated protein 41 (*VPS41*) that was shown to associate with SARS-CoV-2 Orf3 protein^31^ (**Figure S2C**). A very dense cluster of both proviral and antiviral factors belonged to transcriptional regulation and epigenetic modifications networks (p value = 8.08E-9) (**Figure 2** – bottom left, **S2D**), including histone modifiers such as the lysine demethylase KDM1A - also previously identified as a host factor involved in SARS-CoV-2 replication^15^, and regulators of signal transduction such as the *JAK1* tyrosine kinase. Another significant cluster was nicotinate and nicotinamide metabolism (p value = 7.81E-20) (**Figure S2E**), encompassing factors such as the entry receptor for SARS-CoV-2 *ACE2* and one of its regulators, the adipokine Apelin (*APLN*)^14,36,37^. We also observed, as expected, enrichment in pathways involved in the innate immune and antiviral response (p value= 2.35E-11), which were in network with SARS-CoV-2 proteins Orf3, Orf7b and M (**Figure S2F**). Lastly, there was a strong enrichment in factors involved in MAPK signalling (p value = 2.64E-19) (**Figure S2G**), including cell adhesion molecule *CTNNA1*, displayed in our network to interact with SARS-CoV-2 Orf7b protein and to be phosphorylated in response to infection. Overall, these analyses revealed host factors and networks that are supported by one or more OMICs datasets, thus providing a higher level of confidence and more insight into their mechanism of proviral or antiviral action.

**Figure 2.**
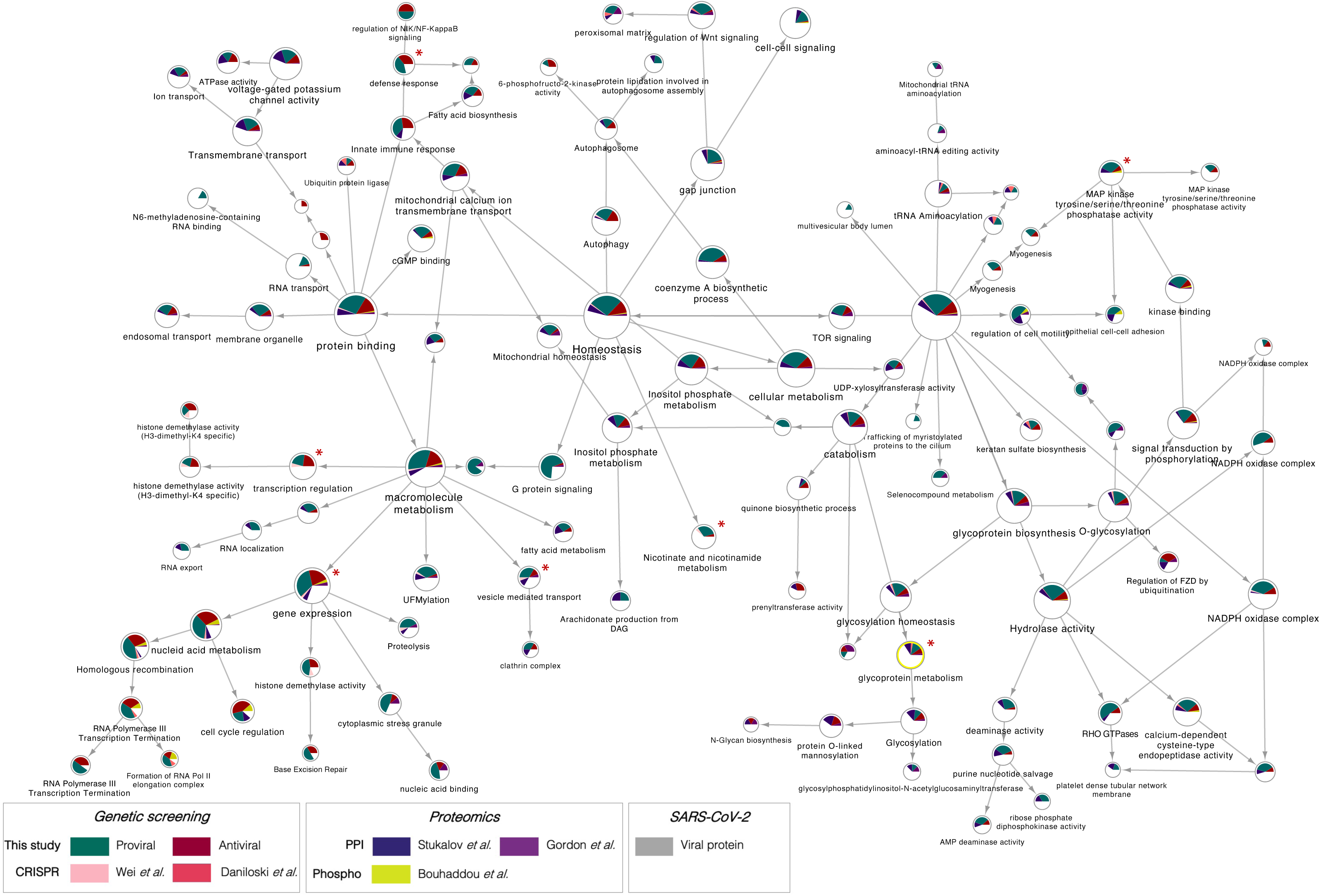
Network integration reveals transcriptional control, epigenetic modifications, and MAPK signalling as relevant networks implicated in SARS-CoV-2 replication. The network containing the identified proviral (**green**) and antiviral (**red**) human host factors was integrated with host factors reported to be relevant for SARS-CoV-2 infection. These include genetic CRISPR screen hits (Wei et al., 2020, **light pink**; Daniloski et al., 2020, **dark pink**), protein-protein interaction hits (Stukalov et al., 2020, **blue**; Gordon et al., 2020, **purple**), as well as hits from a phosphoproteomics study (Bouhaddou et al, 2020, **yellow**). The network was subjected to supervised community detection^66,72^, and the resultant hierarchy is shown. Each node represents a cluster of densely interconnected proteins, and each edge (arrow) denotes containment of one community (edge target) by another (edge source). Labels indicate enriched biological processes. The percentage of each community that corresponds to each dataset is shown by matching colors. Edges indicate interactions from STRING database. **Grey** nodes indicate SARS-CoV-2 proteins. **White** denotes proteins in network (based on STRING) but not identified in any of the OMICs studies. * indicates highlighted clusters.

### Mapping of host factors into SARS-CoV-2 infectious cycle reveals a direct interaction between perlecan and SARS-CoV-2 S protein

The proviral host factors that were found to affect replication of SARS-CoV-2 with two or more siRNAs were evaluated for their effect during the three main stages of the SARS-CoV-2 infectious cycle: entry, replication and assembly/egress. First, to identify host factors involved in viral entry, siRNA-transfected Caco-2 cells were infected with a vesicular stomatitis virus (VSV) encoding luciferase, pseudotyped with either SARS-CoV-2 S protein or VSV Glycoprotein (G), and luciferase levels were measured as indicators of entry. siRNA-mediated knockdown of *ACE2, TMPRSS2, COPB1, ATP6V0C, CLTC, APLN, HSPG2, IRLR2, LIME1* and *AP1G1* significantly reduced entry mediated by SARS-CoV-2 S protein (**Figure 3A**). Of these, *CLTC* and *COPB1* were also found to participate in VSV-G mediated entry (**Figure S3A**), suggesting that both SARS-CoV-2 and VSV hijacked clathrin-mediated endocytosis to enter the host cells. Notably, the other eight factors showed no effect on VSV-G-mediated entry (**Figures 3A, S3A**), including *TMPRSS2* or transmembrane protein *LIME1*, suggesting they are specific for SARS-CoV-2 S-dependent entry.

**Figure 3.**
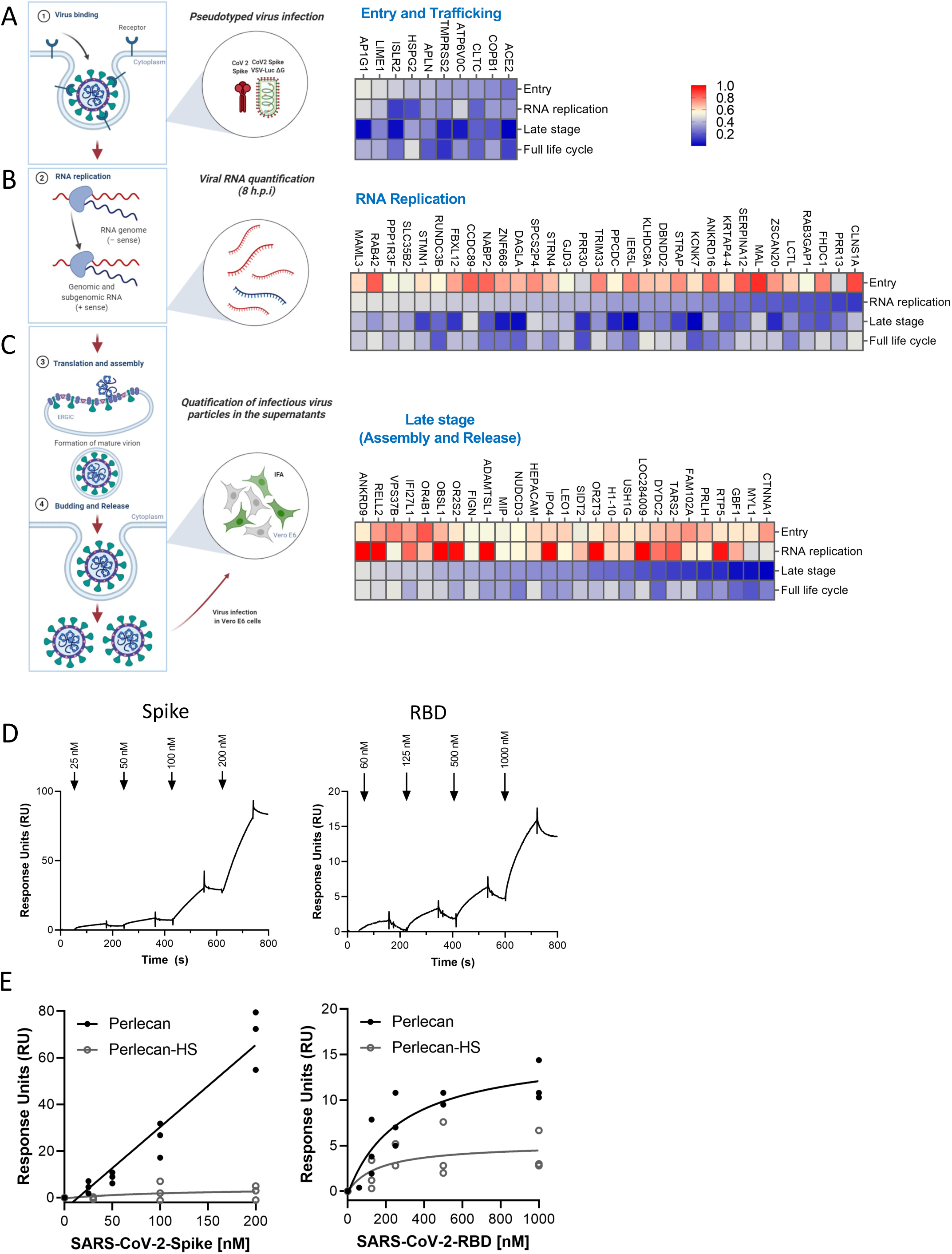
Mapping of host factors into the SARS-CoV-2 replication cycle reveals a direct interaction between entry factor perlecan and SARS-CoV-2 S protein. (A) Caco-2 cells were subjected to siRNA-mediated knockdown of indicated host factors and then infected with SARS-CoV-2 pseudotyped VSV luciferase virus (VSV-S-luc) for 18h prior to measurement of luciferase signal. (B) In parallel, cells were subjected to synchronized infection with SARS-CoV-2 (MOI = 5) for 8h prior to measurement of viral RNA, or (C) supernatants collected at 18h post-infection were used to infect naïve Vero E6 cells. The % of infected cells was then determined at 18h post-infection using immunostaining for viral N protein (3-4). In parallel to these experiments, the impact of depleting these factors on SARS-CoV-2 replication was evaluated at 24 h post-infection in Caco-2 cells (full replication cycle, Figure 3A-C). Results are summarized in the heat map and show the mean (n=2) of relative activities compared to cells treated with non-targeting scramble siRNA. (D and E) Surface plasmon resonance (SPR) was used to evaluate binding of S protein and RBD to perlecan or perlecan without HS spike binding to immunopurified perlecan isolated from human coronary artery endothelial cells. Control flow channels contained immobilized BSA. S protein at indicated concentrations was run across the flow channels for 120 s and dissociation was measured in the following 600 s. The RU values throughout the experiment for BSA were subtracted from the RU values for perlecan to determine the level of specific binding. This experiment was repeated with perlecan treated with heparinase III.

*HSPG2*, also known as Perlecan, was found to be important for SARS-CoV-2 entry (**Figure 3A**). Perlecan is an extracellular proteoglycan, commonly found in all native basement membranes^38^. Heparan sulfate (HS), which is a common modification of Perlecan, has been shown to act as a co-receptor or an attachment factor for a number of viruses, including SARS-CoV-2^39,40^. To test if Perlecan directly interacts with SARS-CoV-2 S protein, we isolated Perlecan from human coronary artery endothelial cells as previously described^41^ and measured its interaction with recombinant full-length S protein and its receptor binding domain (RBD) using a biacore biosensor. Both S and S RBD bound to Perlecan but not albumin (negative control) (**Figure 3D, S3B-C**), although the interaction was more significant with full-length S, illustrated by a higher response units (RU) value (**Figure 3D**). Treatment of the isolated Perlecan with an HSase eliminated binding, showing that the S protein interacts with the HS chain and not the core protein (**Figure 3E**). This is in agreement with previous data showing that HS is required for S binding to cells^40^. Collectively, this data suggests that HSPG2 facilitates SARS-CoV-2 entry and directly interacts with S protein.

Next, to define host factors that affect SARS-CoV-2 RNA replication and translation, viral RNA levels were quantified at 8 h post-infection in Caco-2 cells knockdown for each target gene (**Figure 3B**). This assay revealed 32 host factors that strongly inhibit SARS-CoV-2 RNA replication (>50% inhibition), but have no effect on viral entry. These include RNA-binding protein STRAP, which was previously reported as a SARS-CoV-2 interactor^31^, and the ubiquitin ligase FBXL12, a reported interactor of SARS-CoV-2 Orf8^32^. Lastly, to identify factors involved in the late stages of the viral cycle, we infected naïve Caco-2 cells with viral supernatants that were collected at 18 h post infection of siRNA-transfected Caco-2 cells (**Figure 3C**) followed by immunostaining for viral N protein. We found that depletion of 27 host factors lowered by >50% the amount of infectious viral particle production without affecting viral entry or RNA replication, suggesting that they specifically participate in the late stages of SARS-CoV-2. These include the lysosomal protein SIDT2, which is in agreement with previous reports showing that SARS-CoV-2 hijacks lysosomes for egress^42^, the adhesion molecule CTNNA1, the member of the PAF complex LEO1, shown previously to be targeted by influenza A virus to suppress the antiviral response^43^, and the Golgi resident and vesicle trafficking protein GBF1, a previously reported interactor of SARS-CoV-2 M^31^ (**Figure 3C**).

### Comparative screening reveals potential pan-coronavirus host factors

Motivated by the premise that the identification of host factors essential for replication of several related viruses might inform broad-acting antiviral therapies, we prioritized 47 validated SARS-CoV-2 proviral host factors based on their level of activity, and evaluated their impact on SARS-CoV-1 and MERS replication. From these, 17 factors were required for all three coronaviruses (**Figure 4A**). These include the palmitoyltransferase ZDHHC13, which has been linked to S-mediated syncytia formation and viral entry^44^, the mitochondrial TARS2, a reported interactor of SARS-CoV-2 M protein^32^, and the sorting protein VPS37B, which was previously associated with HIV-1 budding^45^, and was found in our analysis to affect SARS-CoV-2 egress (**Figure 3C**). In addition, eight host factors, including ACE2, AP1G1, and ACE2 positive regulator APLN, whose knockdown reduced ACE2 protein levels^37^ (**Figure 4B**), were required for SARS-CoV-1 and SARS-CoV-2 infection, but had limited effects on MERS-CoV infection. Collectively, these data has revealed a subset of host factors that are conserved across these three coronaviruses and have the potential to lay the groundwork for broad-acting anti-coronavirus therapies.

**Figure 4.**
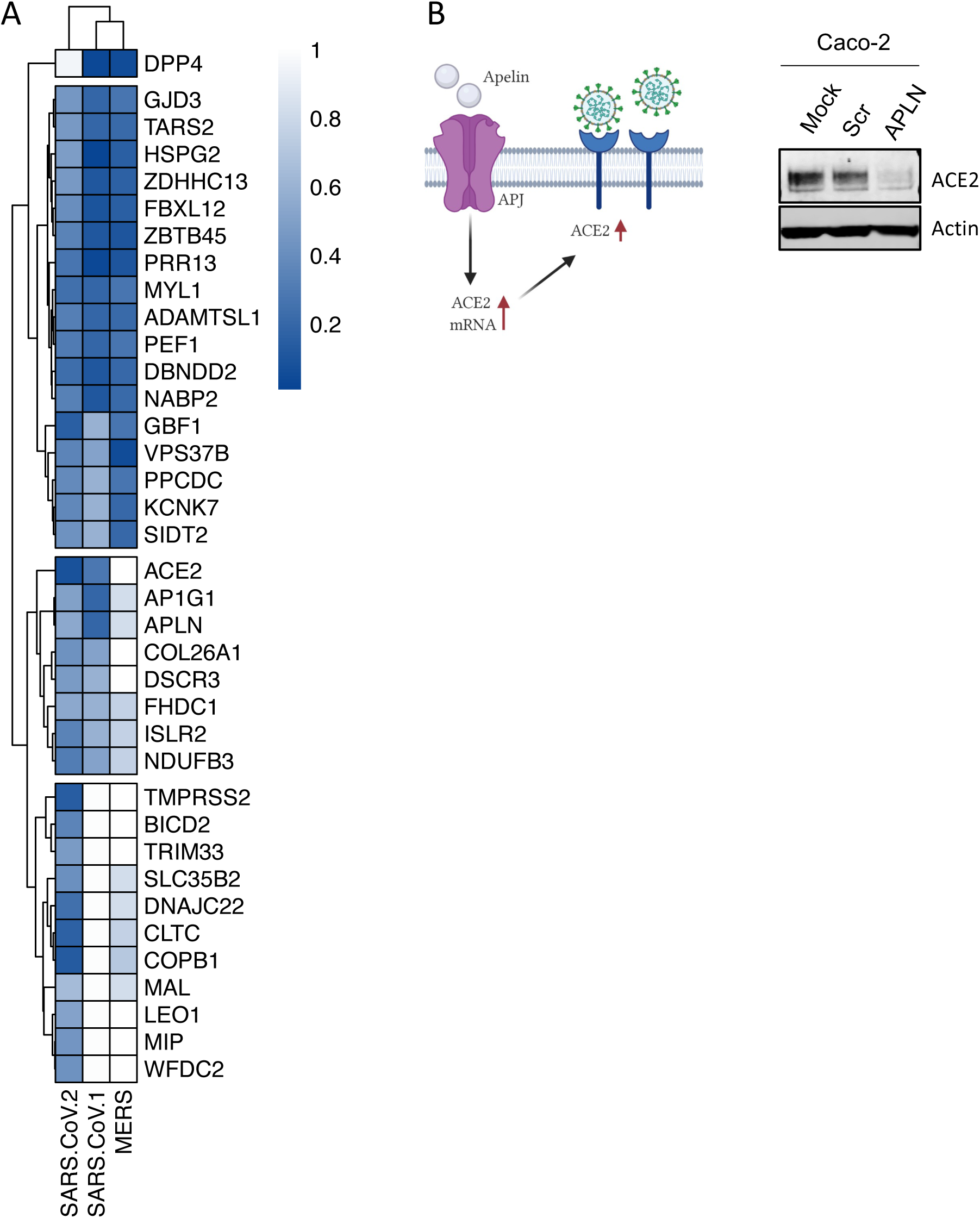
Comparative screening reveals potential pan-coronavirus host factors. (A) Heat map showing normalized infection of SARS-CoV-1, CoV-2, and MERS upon knockdown of indicated human host factors. Caco-2 cells depleted for indicated factors were infected with SARS-CoV-2 (MOI = 0.625) for 48h prior to immunostaining for viral N protein. Shown is quantification of the normalized infection (% of SARS-CoV-2 N^+^ cells) relative to control cells (scrambled siRNA). A549-DPP4 or A549-ACE2 were depleted for indicated factors and then infected with MERS or SARS-CoV-1, respectively (both at MOI 0.1). At 48 h post-infection, supernatants were collected and used to calculate the TCID50. Data shows TCID50/ml relative to control cells (scrambled siRNA). Data show mean ± SD from one representative experiment in duplicate (n=2) of two independent experiments. (B) Cell lysates from Caco-2 cells mock-treated or treated with scrambled or APLN siRNAs for 48 h were then subjected to SDS-PAGE and immunoblotted using antibodies specific for ACE2 and Actin (loading control). Blot is representative of two independent experiments.

### Pharmacological inhibition of BIRC2 reduces SARS-CoV-2 replication *in vitro* and *in vivo*

BIRC2 was one of the proviral host factors identified in our screen (**Table S1**). We previously reported BIRC2 as a critical host factor involved in HIV-1 transcription, through its role as a repressor of the non-canonical NF-κB pathway^46^. Degradation of BIRC2 results in the accumulation of NF-κB- inducing kinase (NIK) and the proteolytic cleavage of p100 into p52, so that p52 can then bind the RELB transcription factor to undergo nuclear translocation and induce the expression of target genes^47^. To evaluate whether pharmacological inhibition of BIRC2 had an impact on SARS-CoV-2 replication, we employed two different BIRC2-specific small molecule antagonists, known as Smac mimetics, AZD5582 and SBI-095329^46,48^. First, we validated the impact of BIRC2 inhibition on NF-κB signalling as treatment of Caco-2 cells with AZD5582 resulted in cleavage of p100 to p52 in a dose-dependent manner (**Figure S4A**). Importantly, we also confirmed that treatment with either AZD5582 or SBI-095329 reduced SARS-CoV-2 infection in a dose-dependent manner without inducing cytotoxicity (**Figure 5A**). To further evaluate the impact of BIRC2 inhibition on SARS-CoV-2 replication *in vivo*, mice were pre-treated with AZD5582 (3 mg/kg), Nirmatrelvir (200 mg/kg), or DMSO (control) and then infected with SARS-CoV-2 (Omicron BA.5 and Alpha B.1.1.7) (**Figure 5B, Figure S4B**). Although prolonged treatment (6 days) with AZD5582 was not well tolerated and resulted in a significant reduction in mice body weight and survival (**Figure S4C-D**), at 3 days post-infection treatment with AZD5582 significantly reduced SARS-CoV-2 viral titers and RNA copy number in the lung both for Omicron and Alpha variants (**Figure 5C-D, Figure S4E**). Combined, these data show that BIRC2 positively impacts SARS-CoV-2 replication *in vitro* and *in vivo*, suggesting its potential as a druggable target for SARS-CoV-2 treatment.

**Figure 5.**
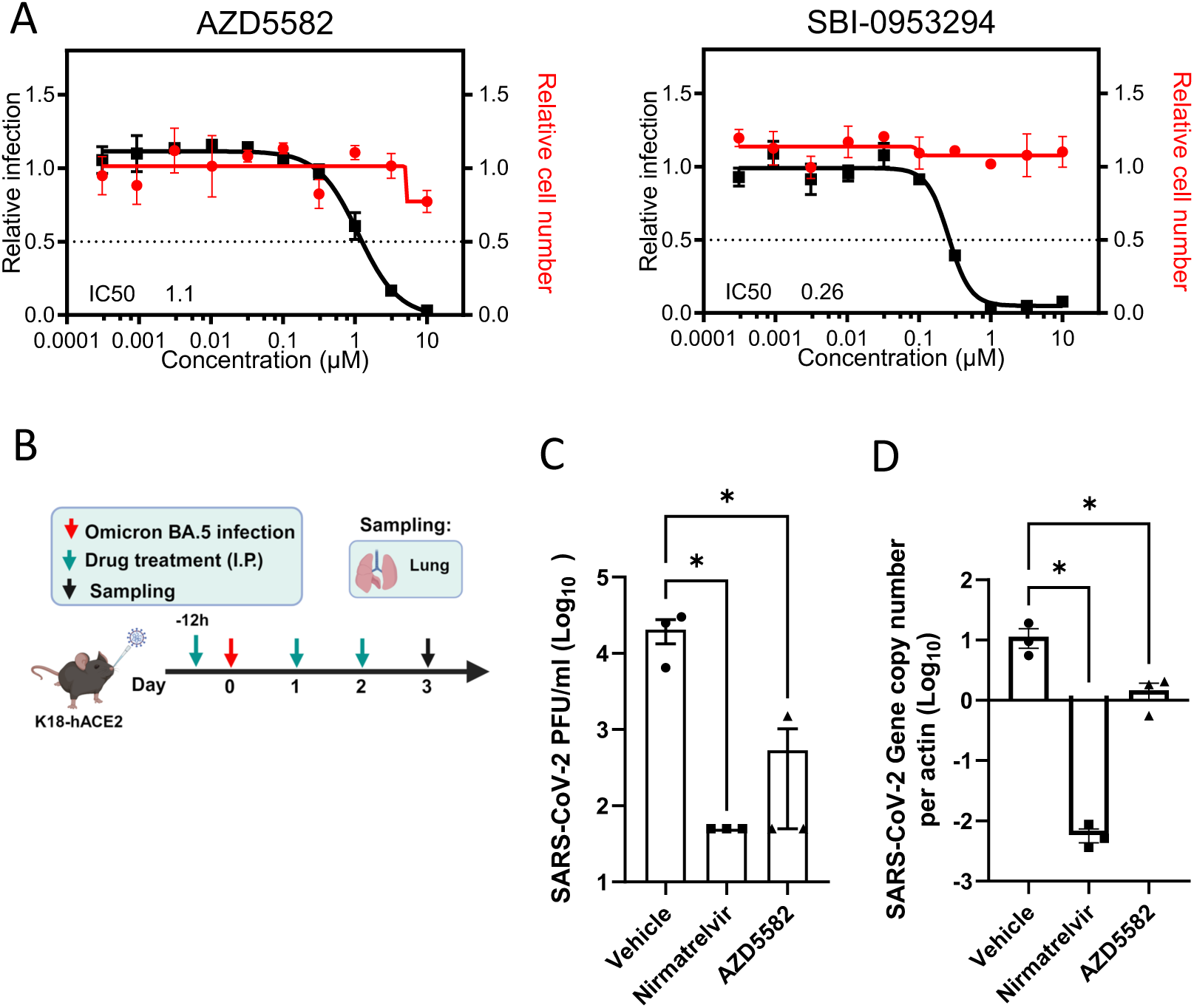
Pharmacological inhibition of BIRC2 reduces SARS-CoV-2 replication *in vitro* and *in vivo*. (A) Dose–response analysis of SBI-0953294 and AZD5582 showing infectivity (black), cell number (red) and cellular IC_50_ values. (B) Layout of mice experiments. Effect of AZD5582 on SARS-CoV-2 Omicron replication in the lungs of infected mice as measured by plaque assay (C) and qRT-PCR (D). Tissue sampling was done at 72hpi. One-way ANOVA when compared with the vehicle control group, *p<0.05. And the detection limit=50 PFU/ml in a 12-well plate.

## DISCUSSION

In this study, we carried out a genome-wide siRNA screen to identify host factors involved throughout the complete SARS-CoV-2 infectious cycle, from attachment and entry to release of viral particles. These data were able to highlight host factors, and networks, supported by multiple OMICs measurements that are required for the replication of SARS-CoV-2 and other coronaviruses, thus constituting relevant therapeutic targets for host-directed antivirals.

Since the beginning of the COVID-19 pandemic, several groups have utilized whole-genome pooled CRISPR screens to identify host factors involved in SARS-CoV-2 replication. Overall, the screens used different cell lines (Vero E6, A549, Huh7.5, Huh7, Calu-3, UM-UC-4, HEK-293), libraries, experimental conditions, and analysis pipelines^15–25^. Comparison of the top hits from some of these pooled screens revealed limited overlap at the gene level, including 91 host factors identified in two or more screens (8.60%), from which 15 were also found in our siRNA screen. GO analysis on these overlapping factors revealed endosomal transport (logP = −9.35686), chromatin remodelling (logP = −7.96025), symbiotic interaction (−logP = −7.01573), vacuole organization (−6.42929), and regulation of DNA methylation (−logP = −6.42929) as the top five enriched biological processes.

Pooled CRISPR screens tend to be biased towards identifying factors that play a role in the early stages of the viral cycle. In contrast, arrayed siRNA screens do not show this bias and capture the entire replication cycle. Accordingly, we found that 40% (4 out of 10) of the siRNA hits assigned to the early steps of the cycle were described in at least one pooled CRISPR screen, while only 6% (2 out of 32) and 4% (1 out of 27) of the hits mapped to replication or the late stages, respectively, were identified as top hits in those screens (**Table S3**). Considering that 85.5% of the host factors identified by the siRNA screen were found to affect post-viral entry stages (**Figure 3**), these data provide novel insights into the poorly understood host factors required for SARS-CoV-2 assembly, trafficking, and budding.

Integration of OMICs datasets can reveal host factors and networks with multiOMIC support thereby increasing the likelihood that they are critical for SARS-CoV-2 replication. In particular, integration of the data generated in this study with a CRISPR functional screen and proteomics - including protein-protein interactions (PPI) and phosphoproteomics - revealed enrichment in four major gene ontology (GO) categories. These are cellular homeostasis, including autophagy or cell-to-cell signalling; gene expression and transcription regulation, including epigenetic regulation and DNA damage; protein binding, including vesicle transport and innate immune regulation; and metabolism, including posttranslational modifications (glycosylation or ubiquitination), and MAPK signalling (**Figure 2**). In fact, several groups have reported critical physical and functional interactions between SARS-CoV-2 and the autophagy machinery to promote viral survival^49,50^, the role of glycosylation to enable S-mediated entry and stimulate innate immune activation^51^, or the ability of SARS-CoV-2 to hijack MAPK11 to promote viral replication^52^. Less understood is the role of epigenetic regulation during SARS-CoV-2. Although it may seem surprising that a cytoplasmic virus relies on nuclear factors to complete its infectious cycle, several cytoplasmic RNA viruses undergo nuclear translocation, are able to mislocalize nuclear proteins into the cytoplasm, or rely on the cytoplasmic products of nuclear transcription factors or associated proteins^53–55^. In addition, recent work showed that SARS-CoV-2 variants of concern have gained the ability to interact with members of the gene transcription regulator PAF complex^56^, including LEO1, which was found as a validated host factor in our screen (**Figure 1E**). However, more work will be required to understand the functional consequences of these interactions and mechanism of action.

Among the factors found to affect SARS-CoV-2 entry was HSPG2 (Perlecan, **Figure 3A**). Perlecan is a large, multi-domain proteoglycan modified by HS that is located in the extracellular matrix (ECM) and basement membranes of the airway and alveolar epithelia and could therefore directly abet SARS-CoV-2 infection^38^. Subsequently, we employed Surface Plasmon Resonance (SPR) and revealed Perlecan as a direct interactor of SARS-CoV-2 S protein, thus adding to the growing evidence that HS-modified proteins could participate in SARS-CoV-2 entry. Studies utilizing enzymatic degradation of HS or using competitive inhibitors that block the binding sites of HS have demonstrated reduced infection rates of SARS-CoV-2 in cell cultures^57^. Furthermore, variations in the structure of HS chains can affect the efficiency of viral attachment and entry, indicating a level of specificity in the interaction between HS and SARS-CoV-2. The involvement of HS in the entry mechanism of SARS-CoV-2 is also consistent with their known roles in the entry of other viruses^58^. Further understanding of this mechanism could lead to broad-spectrum antiviral strategies targeting the initial attachment phase of viral infection. Another potential mechanism of broad-acting viral inhibition is targeting the inhibitor of apoptosis proteins (IAP), which play key and complex roles in innate immunity, inflammation as well as the regulation of cell death and cell proliferation^59,60^. Smac mimetics inhibit IAPs and have been recognized as potent HIV-1 latency reversal agents^46^, and more recently described to have antiviral properties^48^. In this study, we found two Smac mimetics, AZD5582 and SBI-095329, that through inhibition of the proviral host factor BIRC2, conferred antiviral properties *in vitro* against the ancestral Wuhan-1 SARS-CoV-2, and *in vivo* (AZD5582) across the two variants of concern Omicron and Alpha. Although no toxicity was recorded in our *in vitro* experiments, prolonged treatment in mice resulted in reduced survival and body weight, suggesting more work will be required to address their safety profile. Importantly, a very recent publication showed that the Boehringer Ingelheim Smac mimetic BI-82, which is orally available, conferred antiviral activities across dengue, zika, and hepatitis B virus (HBV) *in vitro*, and was well-tolerated and showed potent efficacy against influenza A virus *in vivo*^48^. Combined with our data, this suggests that the expression program governed by non-canonical NF-1B signalling potently restricts SARS-Cov-2 replication both *in vitro* and *in vivo*, and further underscore the potential of Smac mimetics as broad-acting antiviral therapies.

In summary, our study unveils novel host factors that are critical for all three main stages of SARS-CoV-2 infectious cycle. Importantly, we carried out comparative screening across SARS-CoV-1 and MERS highlighting commonalities that could inform the development of host-directed, pan-coronaviral antiviral therapies.

## METHODS

### Cells and Viruses

SARS-CoV-2 USA-WA1/2020, isolated from an oropharyngeal swab from a patient with a respiratory illness who developed clinical disease (COVID-19) in January 2020 in Washington, USA, was obtained from BEI Resources (NR-52281). These viruses were propagated using Vero E6 cells, collected, aliquoted, and stored at −80 °C. Plaque forming unit (PFU) assays were performed to titrate the cultured virus. All experiments involving live SARS-CoV-2 followed the approved standard operating procedures of the Biosafety Level 3 facility at the Sanford Burnham Prebys Medical Discovery Institute. SARS-CoV-1 (MA15) was generated produced as decribed^61^. The Jordan MERS-CoV strain (GenBank accession no. KC776174.1, MERS-CoV-Hu/Jordan-N3/2012) was kindly provided by Kanta Subbarao (National Institutes of Health, Bethesda, MD) and Gabriel Defang (Naval Medical Research Unit-3, Cairo, Egypt). All work with SARS-CoV-1 and MERS was performed in a Biosafety Level 3 laboratory and approved by the University of Maryland Institutional Biosafety Committee. Caco-2 (ATCC HTB-37), Vero E6 (ATCC CRL-1586), HEK293T (ATCC CRL-3216), Calu-3 (ATCC HTB-55), A549-DPP4 (kind gift from Susan Weiss, UPenn), and A549-ACE2 (kind gift from Brad Rosenburg, Mount Sinai) cells were maintained in cell growth media: Dulbecco’s modified eagle medium (DMEM, Gibco) supplemented with 10 % heat-inactivated fetal bovine serum (FBS, Gibco), 50 U/mL penicillin −50 µg/mL streptomycin (Fisher Scientific), 1 mM sodium pyruvate (Gibco), 10 mM 4-(2-hydroxyethyl)-1-piperazineethanesulfonic acid (HEPES, Gibco), and 1X MEM non-essential amino acids solution (Gibco). All cells were regularly tested and were confirmed to be free of mycoplasma contamination.

### siRNA screening

A whole-genome wide ON-TARGETplus SMARTpool siRNA library (Dharmacon, each containing 4 siRNAs targeting an individual gene) was seeded at 0.5 pmol each/well in 384-well plates (Greiner). For reverse transfection, Lipofectamine RNAiMAX was added in 10 1L OPTI-MEM to each well at a final dilution of 1:100 using a Combi reagent dispenser, followed by addition of 3,000 Caco-2 cells in 40 1L complete media per well. 48h post transfection, cells were challenged by SARS-CoV-2 at MOI 0.625. 48h post infection, plates were fixed by 4% PFA in PBS for 4h at room temperature, then permeabilized by 0.4% Triton X-100 in PBS for 15min at room temperature. Plates were blocked by 10% goat serum in 3% BSA in PBS for 30min at room temperature, followed by incubation of primary antibody against SARS-CoV-2 NP at 1,000 in 3% BSA in PBS at 4°C overnight. Primary antibody inoculum was removed and plates were washed 3 times with PBS by plate washer, then incubated with anti-rabbit Alexa Fluor 488 (Invitrogen) at 1,000 in PBS for 1h at room temperature. Secondary antibody inoculum was removed and plates were washed 3 times with PBS by plate washer, then DAPI was added in PBS. Plates were then sealed and imaged using the Celigo Image Cytometer (Nexcelom)..

### Generation of Calu-3 CRISPR/Cas9 knockouts

Detailed protocols for RNP production have been previously published^62^. Briefly, lyophilized guide RNA (gRNA) and tracrRNA (Dharmacon) were suspended at a concentration of 160 µM in 10 mM Tris-HCL, 150mM KCl, pH 7.4. 5µL of 160µM gRNA was mixed with 5µL of 160µM tracrRNA and incubated for 30 min at 37°C. The gRNA:tracrRNA complexes were then mixed gently with 10µL of 40µM Cas9 (UC-Berkeley Macrolab) to form CRISPR-Cas9 ribonucleoproteins (crRNPs). Five 3.5µL aliquots were frozen in Lo-Bind 96-well V-bottom plates (E&K Scientific) at −80°C until use. Each gene was targeted by 4 pooled gRNA derived from the Dharmacon pre-designed Edit-R library for gene knock-out (sequences and catalog numbers provided in the table below). Non-targeting negative control gRNA (Dharmacon, U-007501) was delivered in parallel. Each electroporation reaction consisted of 2.0x10^5 Calu-3 cells, 3.5 µL crRNPs, and 20 µL electroporation buffer. Calu-3 cells were grown in fully supplemented MEM (10% FBS, 1xPen/Strep, 1x non-essential amino acids) to 70% confluency, suspended and counted. crRNPs were thawed and allowed to come to room-temperature. Immediately prior to electroporation, cells were centrifuged at 400xg for 3 minutes, supernatant was removed by aspiration, and the pellet was resuspended in 20 µL of room-temperature SE electroporation buffer plus supplement (Lonza) per reaction. 20 µL of cell suspension was then gently mixed with each crRNP and aliquoted into a 96-well electroporation cuvette for nucleofection with the 4-D Nucleofector X-Unit (Lonza) using pulse code EO-120. Immediately after electroporation, 80 µL of pre-warmed media was added to each well and cells were allowed to rest for 30 minutes in a 37°C cell culture incubator. Cells were subsequently moved to 12-well flat-bottomed culture plates pre-filled with 500 µL pre-warmed media. Cells were cultured at 37°C / 5% CO2 in a dark, humidified cell culture incubator for 4 days to allow for gene knock-out and protein clearance prior to downstream applications.

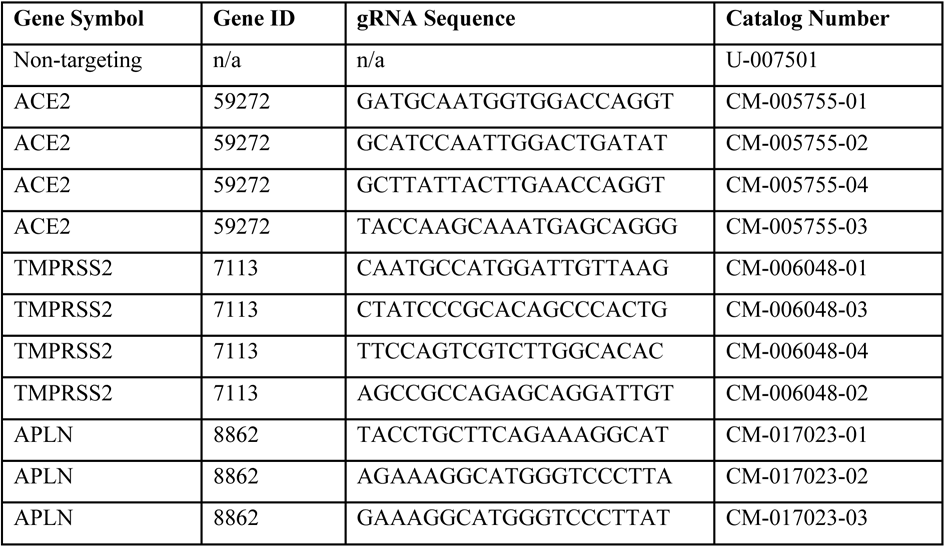
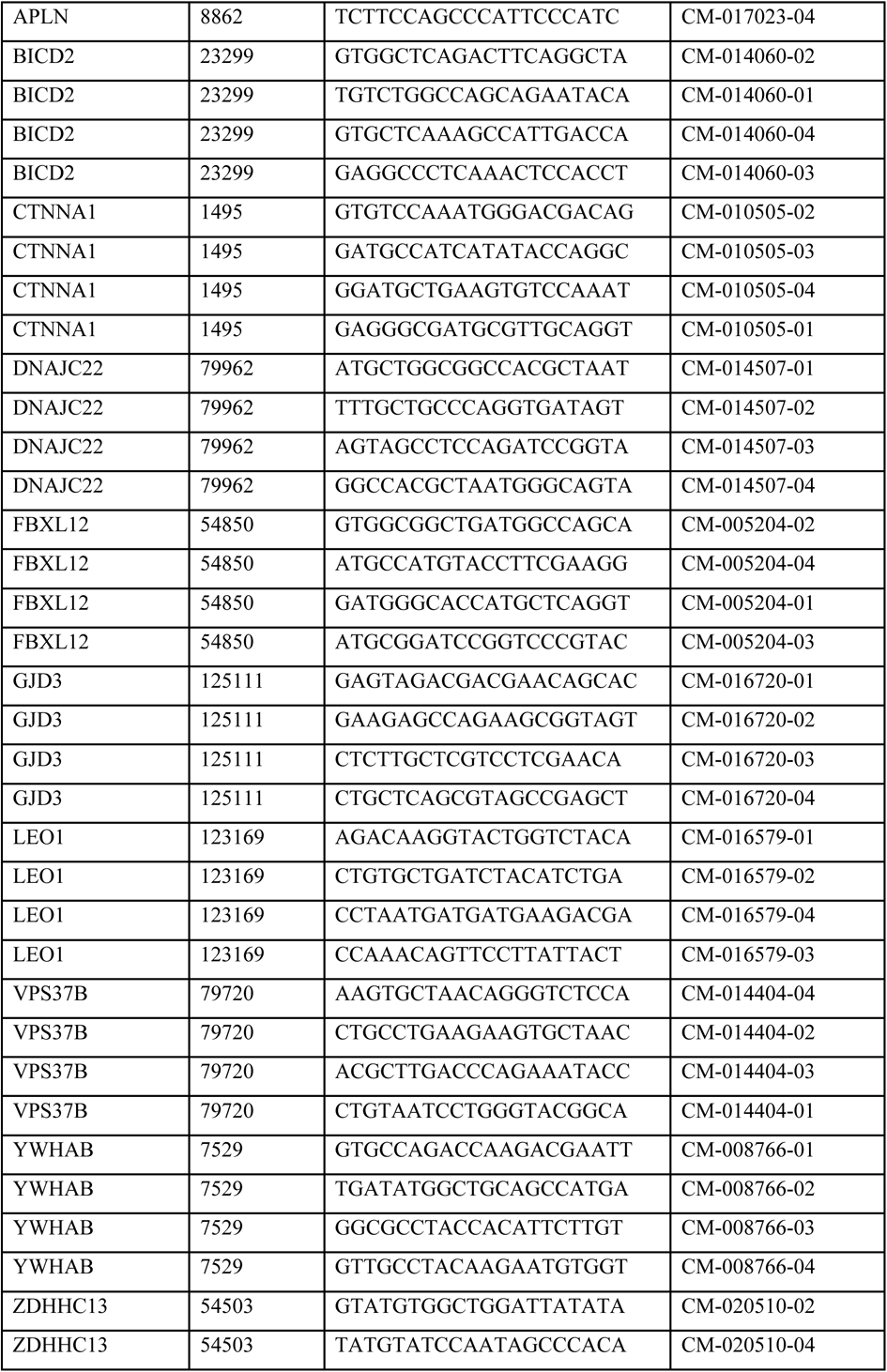
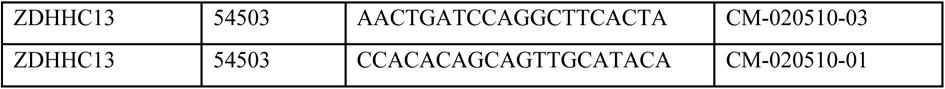

### Network analyses

*Rationale:* To understand the biochemical and functional context in which the identified host factors for SARS-CoV-2 function, we built a model that places these hits in known interactomes. A hierarchy of the clusters is generated wherein larger clusters are composed of smaller ones ^63,64^. Unlike the human-curated Gene Ontology (GO), the structure is derived by the use of a multi-scale clustering algorithm applied to a reference protein-protein interaction network, in this case, a high-confidence subset of the STRING database. To focus the model on the experimental data, it is built using the functional hits found in this study and their close neighbors. The interpretation of the experiment is performed by projecting the hits onto the clusters in the model, analogous to mapping them to GO terms^65^. Candidate names are proposed for each cluster by performing functional enrichment, finding the closest matching pathways and GO terms. Comparing this model to the result of a GO analysis, it has the advantages that its terms (clusters) are algorithmically derived from protein interactions that are in a sense “proximal” to the hits so that the hits can be investigated in the context of their underlying interactions. *Approach:* To explore the highest confidence interactions of “hit” proteins, we selected the STRING - Human Protein Links - High Confidence (Score >= 0.7) protein-protein interaction network available on NDEx as the “background” network (link provided below). We then performed network propagation to select a neighborhood of 300 proteins ranked highest by the algorithm with respect to these seeds ^66^. This “neighborhood” network was extracted from the background network. We then identified densely interconnected regions, i.e. “communities” within the neighborhood network, using the community detection algorithm HiDeF via the Community Detection Application and Service (CDAPS) ^67,68^ (app available at ^24,25^). The result of HiDeF from CDAPS was a “hierarchy” network where each node represented a community of proteins, and edges denoted containment of one community (the “child”) by another (the “parent”). Finally, the hierarchy network was styled, communities were labelled by functional enrichment using gProfiler (via CDAPS), p values were calculated based on the accumulative hypergeometric distribution, and a layout was applied. The STRING - Human Protein Links - High Confidence (Score > = 0.7) network is available in the Network Data Exchange (NDEx) at http://ndexbio.org/#/network/275bd84e-3d18-11e8-a935-0ac135e8bacf.

### Generation pseudotyped SARS-CoV-2 virus

VSV pseudotyped with Spike (S) protein of SARS-CoV-2 wild-type (WT) (Wuhan-Hu-1) were generated according to a published protocol^69^. Briefly, BHK-21/WI-2 cells (Kerafast, MA) transfected with SARS-CoV-2 S protein were inoculated with VSV-G pseudotyped ΔG-luciferase VSV (Kerafast, MA). After a 2h incubation at 37 °C, the inoculum was removed and cells were treated with DMEM supplemented with 5% FBS, 50 U/mL penicillin, and 50 µg/mL streptomycin. Pseudotyped particles were collected 24h post-inoculation, then centrifuged at 1,000×g to remove cell debris and stored at −80°C until use.

### Mapping factors into the SARS-CoV-2 replication cycle

Caco-2 cells were transfected with indicated siRNAs and incubated for 48 h at 37°C, 5% CO2. To determine the effect of the identified factors on viral entry, cells were infected with VSV-S-luciferase or VSV-G-luciferase and incubated for 16h. The activity of firefly luciferase was then quantified using the bright-Glo™ luciferase assay (Promega). To measure RNA replication and late stages, cells were infected with SARS-CoV-2 (USA-WA1/2020) at a MOI 0.625 for 1h on ice. Viral inoculum was removed and cells were washed twice with 1xPBS and supplemented with cell growth media. At 6h post-infection, intracellular viral RNA was purified from infected cells using the TurboCapture mRNA Kit (Qiagen) in accordance with the manufacturer’s instructions. The purified RNA was subjected to first-strand cDNA synthesis using the high-capacity cDNA reverse transcription kit (Applied Biosystems, Inc). Real-time quantitative PCR (RT-qPCR) analysis was then performed using TaqPath one-step RT-qPCR Master Mix (Applied Biosystems, Inc) and ActinB CTRL Mix (Applied Biosystems, Inc) for housekeeping genes, and the following primers and probe for qPCR measurements of viral genes: N-Fwd: 5’-TTACAAACATTGGCCGCAAA-3’; N-Rev: 5’-GCGCGACATTCCGAAGAA-3’; N-Probe: 5’-FAM-ACAATTTGCCCCCAGCGCTTCAG-BHQ-3’. To evaluate late stages, supernatants collected at 18h post-infection were used to infect naïve Vero E6 cells. At 18h post-infection, cells were fixed with 5% PFA (Boston BioProducts) for 4h at room temperature and then subjected to immunostaining and imaging for SARS-CoV-2 N protein.

### Binding of Spike protein to Perlecan

Immunopurified Perlecan isolated from human coronary artery endothelial cells^41^ (10 µg/mL in Dulbecco’s phosphate buffered saline (DPBS) pH 7.4) was immobilized onto gold sensor chips (Sensor chip Au, Cytiva) at 5 µL/min in an SPR system (Biacore T200, Cytiva) at 25 °C for 240s. The sensor chip flow channels were then washed with DPBS at 5 μL/min until a stable response unit (RU) was achieved. The flow channels were then exposed to bovine serum albumin (BSA; 10 mg/mL in DPBS) at a flow rate of 5 μL/min for 240s and washed with DPBS until a stable RU was observed. Control flow channels contained immobilized BSA. Spike protein (25, 50, 100 and 200 nM in DPBS) was exposed to the flow channels at a flow rate of 10 μL/min for 120s. The dissociation of Spike protein was measured in the following 600s. The RU values throughout the experiment for BSA were subtracted from the RU values for Perlecan to determine the level of specific binding. This experiment was repeated with Perlecan treated with heparinase III (0.01 U/mL in DPBS for 16 h at 37 °C; EC 4.2.2.8; Iduron, Cheshire, UK) to remove heparan sulfate (HS). n=3 per condition.

### Evaluation of host factors using SARS-CoV-1 and MERS

A549 cells stably expressing DPP4 or ACE2 were subject to siRNA mediated knockdown of select host factors for 72 hours prior to use. Transfection was performed as described in^70^, modified for a 96 well plate format. A549-DPP4 cells were infected with MERS-CoV (Jordan strain) and A549-ACE2 cells were infected with SARS-CoV (MA15 strain), both at MOI 0.1. 48-hour post infection, supernatant from infected cells was collected and virus titer determined by TCID50 assay (as described^71^). Two experiments were performed and the average TCID50/ml calculated. Scrambled siRNA sequences acted as a negative control and ACE2 and DPP4 targeting siRNAs were positive controls.

### Inhibition of SARS-CoV-2 replication in vitro by Smac mimetics

Caco-2 cells were treated with the compounds (AZD5582 and SBI-0953294) for 18h prior to infection with SARS-CoV-2 (Wuhan-1 isolate) at MOI of 0.625. 48 hours post-infection, the infected cells were fixed with 4% paraformaldehyde for 2 h and permeabilized with 0.5% Triton X-100 for 15min. After blocking with 3% bovine serum albumin (BSA) for 15 min, cells were incubated with rabbit anti-SARS-CoV-2 NP antibodies for 1hours. After two washes with phosphate-buffered saline (PBS), the cells were incubated with Alexa Fluor 488-conjugated goat-anti-rabbit IgG (Thermo Fisher Scientific) for 1 h at room temperature. After two additional washes, the cells were mounted with DAPI (BioLegend) and images were acquired using the Celigo Image Cytometer (Nexcelom).

### *In vivo* experiments

Male K18-hACE2 mice, aged 6-10 weeks old, were kept in biosafety level housing and given access to standard pellet feed and water ad libitum as we previously described. Mice were randomly allocated to experimental groups (n=3 for Omicron experiment, n=11 for Alpha experiment) for antiviral evaluation. All experimental protocols were approved by the Animal Ethics Committee in the University of Hong Kong (CULATR) and were performed according to the standard operating procedures of the biosafety level 3 animal facilities (Reference code: CULATR 5754-21). The experiments were not blinded. Experimentally, each mouse was intranasally inoculated with 10,000 PFU of SARS-CoV-2 (Omicron BA.5) or 200 PFU (Alpha B.1.1.7) in 20 µL PBS under intraperitoneal ketamine and xylazine anaesthesia. Twelve-hours before-virus-challenge, mice were intraperitoneally given either Nirmatrelvir (200 mg/kg), or AZD5582(3 mg/kg) or 1% DMSO in PBS (vehicle control). The second and third doses of drug treatment was performed at 12 and 36 hpi, respectively. For Omicron experiments, three animals in each group were sacrificed at 3dpi for virological analyses (Omicron). Lung tissue samples were collected. Viral yield in the tissue homogenates were detected by plaque assay. For Alpha experiments, animals (n=5) were monitored twice daily for clinical signs of disease. Their body weight and survival were monitored for 14 days or until death. Six animals in each group were sacrificed at 3dpi for virological analyses. Lung tissue samples were collected. Viral yield in the tissue homogenates were detected by plaque assay. A 30% body weight loss is set as human endpoint.

## ETHICS STATEMENT

All experimental protocols with mice were approved by the Animal Ethics Committee in the University of Hong Kong (CULATR) and were performed according to the standard operating procedures of the biosafety level 3 animal facilities (Reference code: CULATR 5754-21).

## DATA AVAILABILITY

The genome-wide siRNA screen data generated in this study have been deposited to Figshare (https://figshare.com/s/4117ac39b1d21b56f5e6).

## STATISTICS

Statistical parameters including the exact value of n, dispersion, and precision measures (mean ± SD or SEM), and statistical significance are reported in the figures and figure legends. Statistical significance between groups was determined using GraphPad Prism v8.0 (GraphPad, San Diego, CA), and the test used is indicated in the figure legends.

## ACKNOWLEDGMENTS

We would like to thank Dr. Tanya Dragic for her insightful comments into the manuscript. We also would like to thank Sylvie Blondelle and Larry Adelman for biosafety support, and Rowland Eaden for shipping assistance. We also would like to thank Kanta Subbarao and Gabriel Defang for the Jordan MERS-CoV strain, Susan Weiss for the A549-DPP4 cells, and Brad Rosenberg for the A549-ACE2 cells. This work was supported by the following grants to the Scripps Research Institute, Sanford Burnham Prebys Medical Discovery Institute, the Icahn School of medicine at Mount Sinai and the University of Hong Kong: DHIPC: U19 AI118610; Fluomics/NOSI-SYBIL: U19 AI135972; HMRF Fellowship: 07210107. This work was also supported by generous philanthropic donations from Dinah Ruch and Susan & James Blair, from the JPB Foundation, the Open Philanthropy Project (research grant 2020-215611 (5384)), a generous grant from the James B. Pendleton Charitable Trust, and anonymous donors. The funding sources had no role in the study design, data collection, analysis, interpretation, or writing of the report.

## AUTHOR CONTRIBUTIONS STATEMENT

L.M.-S., X.Y. and S.K.C., conceived and designed the experiments. L.M.-S., X.Y., Y.P., S.Y., D.F., S.W., L.R., P.D.J., L.M.S., W.J.C., and J.F.H. conducted and/or analyzed experiments. L.M.-S., C.C., D.P., and T.I. generated the network model. S.Y. carried out the animal experiments. L.M.-S., X.Y. and L.P. was responsible for the data visualization and curation. L.M.-S., X.Y. N.M. T.D. and S.K.C. wrote the manuscript with contributions from all authors. Funding Acquisition, J.F.H., M.B.F., T.I., A.G.-S., and S.K.C.

## COMPETING INTERESTS STATEMENT

J.F.H. has received research support, paid to Northwestern University, from Gilead Sciences, and is a paid consultant for Merck. The A.G.-S. laboratory has received research support from GSK, Pfizer, Senhwa Biosciences, Kenall Manufacturing, Blade Therapeutics, Avimex, Johnson & Johnson, Dynavax, 7Hills Pharma, Pharmamar, ImmunityBio, Accurius, Nanocomposix, Hexamer, N-fold LLC, Model Medicines, Atea Pharma, Applied Biological Laboratories and Merck, outside of the reported work. A.G.-S. has consulting agreements for the following companies involving cash and/or stock: Castlevax, Amovir, Vivaldi Biosciences, Contrafect, 7Hills Pharma, Avimex, Pagoda, Accurius, Esperovax, Applied Biological Laboratories, Pharmamar, CureLab Oncology, CureLab Veterinary, Synairgen, Paratus, Pfizer and Prosetta, outside of the reported work. A.G.-S. has been an invited speaker in meeting events organized by Seqirus, Janssen, Abbott, Astrazeneca and Novavax. A.G.-S. is inventor on patents and patent applications on the use of antivirals and vaccines for the treatment and prevention of virus infections and cancer, owned by the Icahn School of Medicine at Mount Sinai, New York, outside of the reported work. All other authors declare no competing interests.

## SUPPLEMENTAL FIGURE LEGENDS

**Figure S1.**
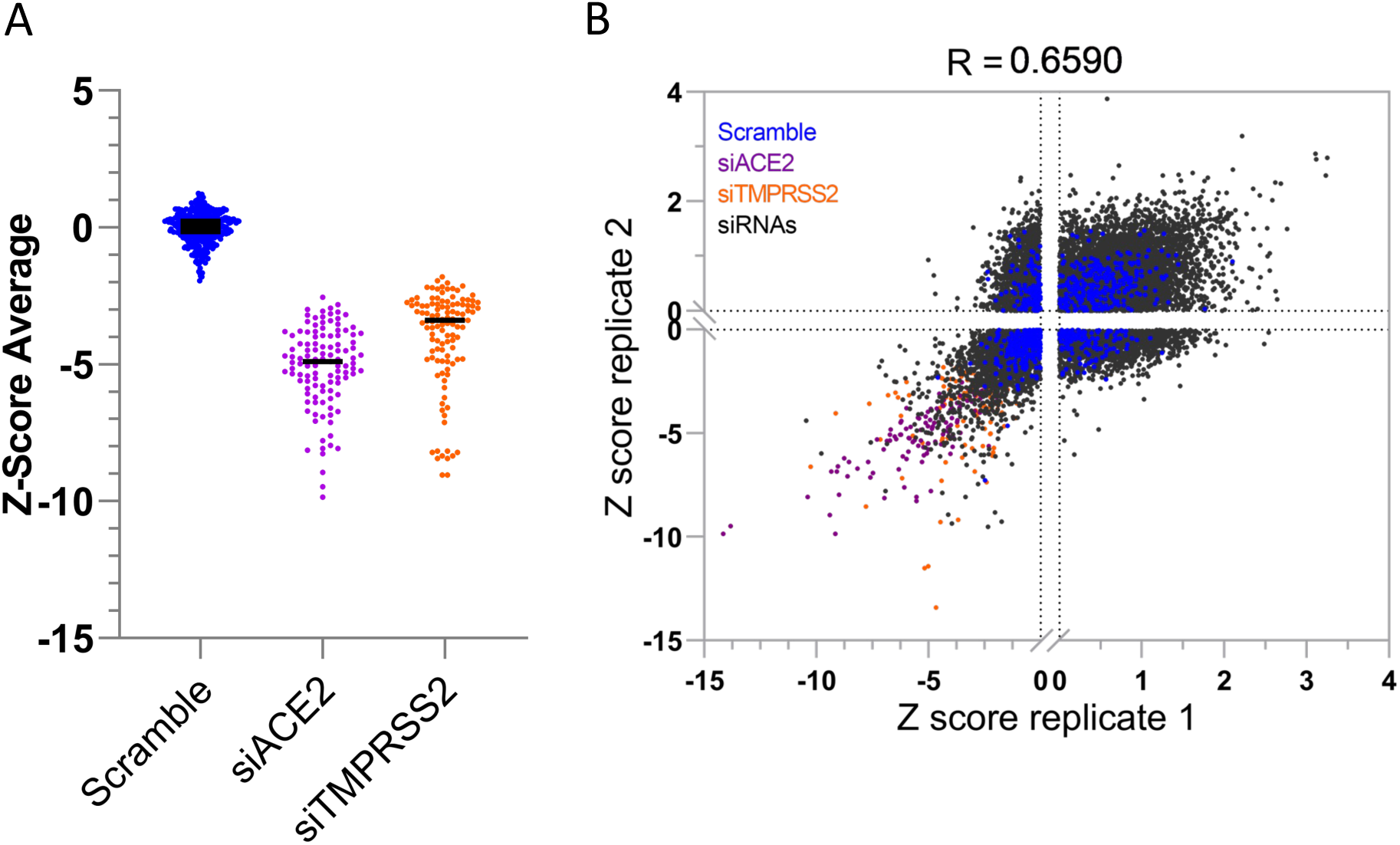
Genome-wide siRNA screen identifies host factors involved in SARS-CoV-2 replication. (A) Dot plot shows average SARS-CoV-2 infectivity Z-score values from the genome-wide siRNA screen. Controls are shown (non-targeting scrambled siRNA, negative; siACE2 and siTMPRSS2, positive). (B) Correlation plots of Z-score values for genome-wide siRNA screens using Caco-2 cells infected with SARS-CoV-2. R = Pearson correlation coefficient between screens.

**Figure S2.**
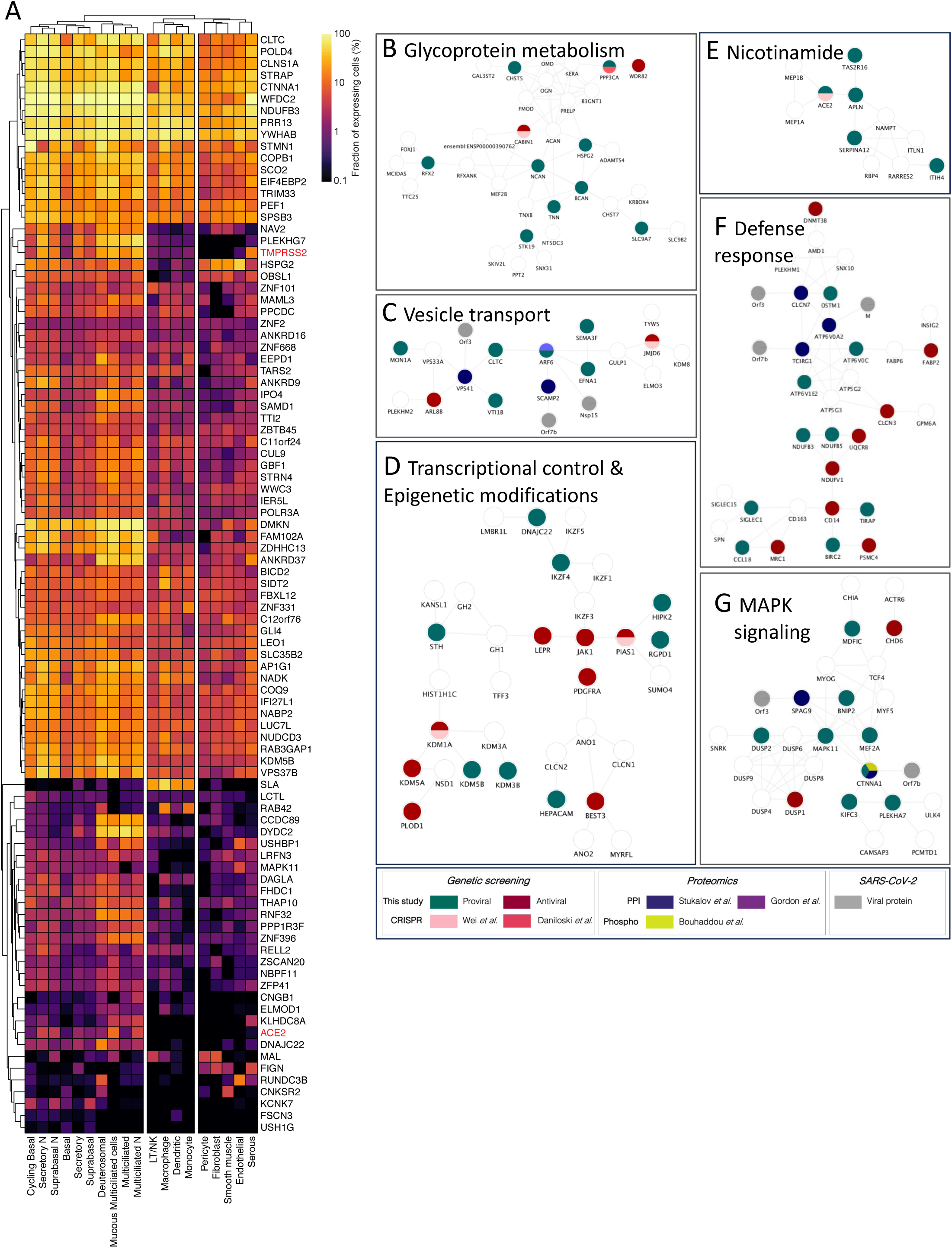
Expression of the identified host factors in SARS-CoV-2 target cells. (A) Heatmap shows percentage of detectable levels of expression of a given factor in the indicated cell type^73^. % expression >1 was considered a detectable level. (B-G) Zoom-in insets from selected biological processes are indicated with an asterisk * in the hierarchy. The nodes indicate host factors and their color matches the dataset where they were identified. Edges indicate interactions from STRING database. Grey nodes indicate SARS-CoV-2 proteins.

**Figure S3.**
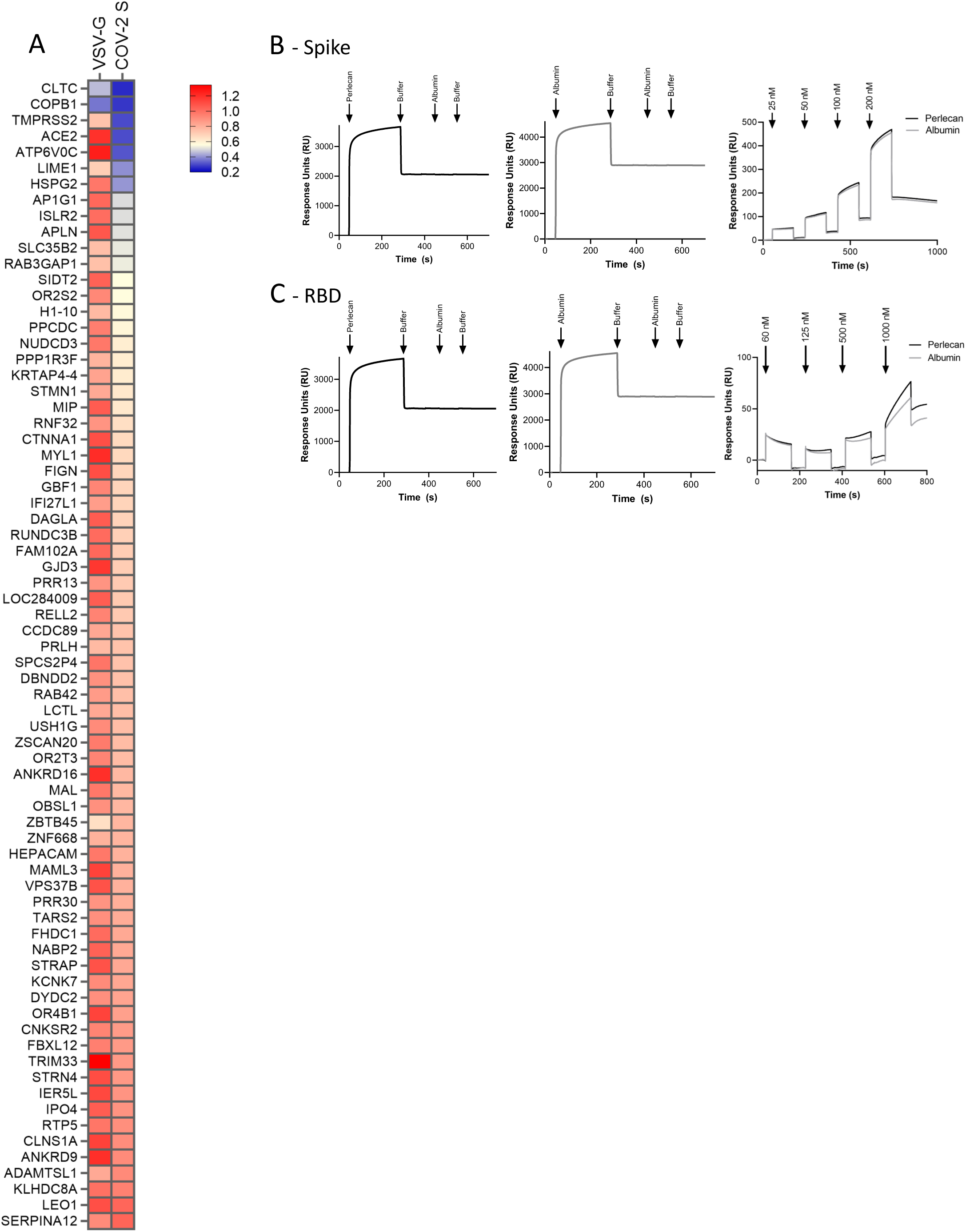
Mapping of host factors into SARS-CoV-2 infectious cycle. (A) Caco-2 cells subjected to siRNA-mediated knockdown of the indicated host factors were infected with SARS-CoV-2 pseudotyped VSV luciferase virus (VSV-S) or VSV luciferase virus expressing its natural glycoprotein (VSV-G) for 18h prior to measurement of luciferase signal. Data represent mean from one representative experiment in duplicate (n=2). (B,C) Binding of spike protein and RBD to perlecan. Surface plasmon resonance (SPR) was used to evaluate spike binding to perlecan. This experiment was repeated twice.

**Figure S4.**
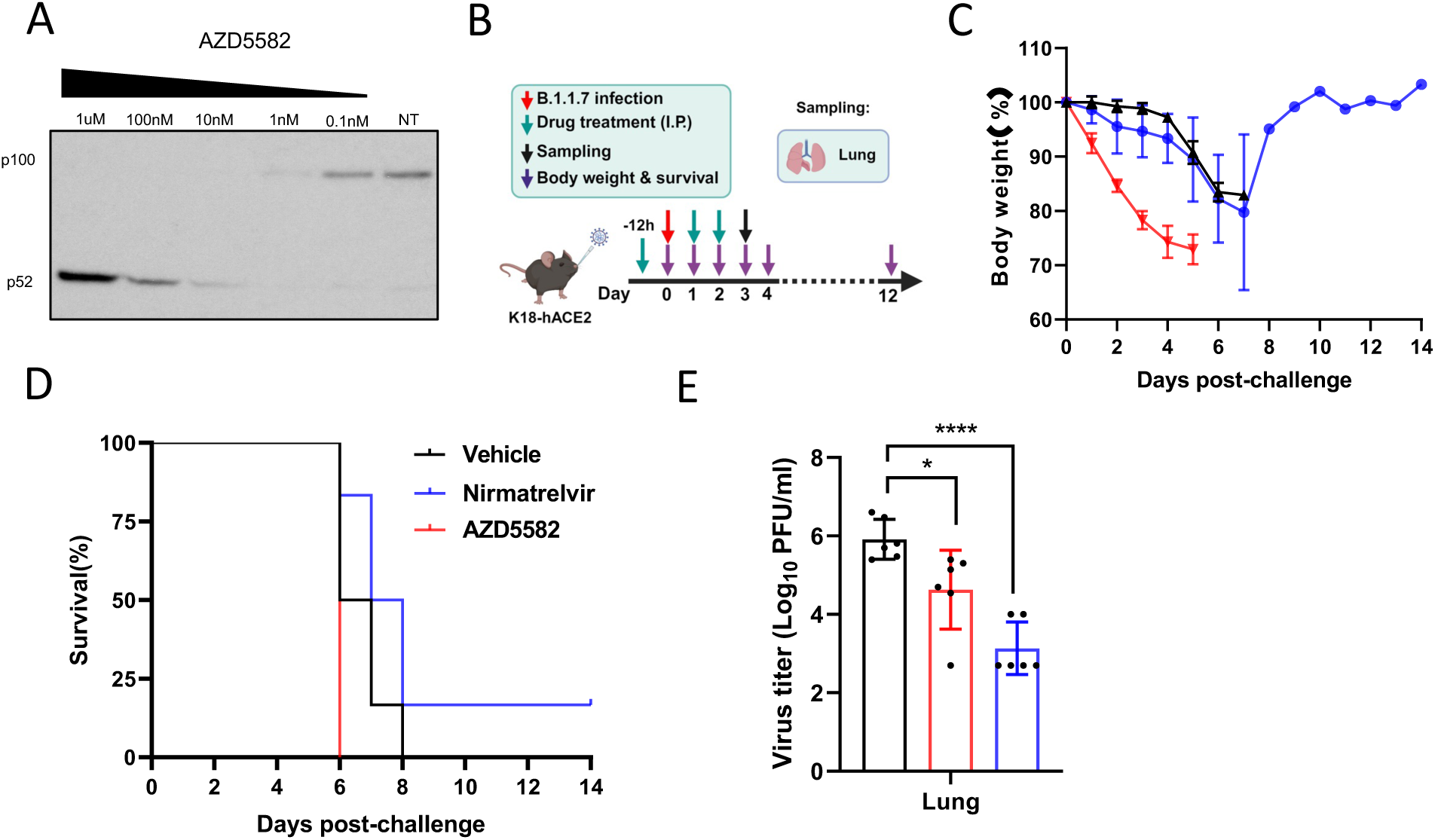
Pharmacological inhibition of BIRC2 reduces SARS-CoV-2 replication *in vitro* and *in vivo*. (A) Cells were treated with AZD5582 at the indicated concentrations. 24 hours post-treatment, the cell lysates were analyzed by Western blotting for p100/p52 protein. A representative immunoblot presented here demonstrate that AZD5582 treatment induces the cleavage of p100. (B) Layout of mice experiments using SARS-CoV-2 B.1.1.7 (Alpha) infection. Effect of AZD5582 on SARS-CoV-2 replication in survival (C) and body weight (D) were recorded for 14 days post-infection. Virus titer as measured in the lungs of infected mice by plaque assay (E) were performed on 3dpi. Tissue sampling was done at 72hpi. One-way ANOVA when compared with the vehicle control group. *P<0.05, ****P<0.001.

